# The genitals of rhinoceros beetles: a general overview of the endophallus in the tribe Agaocephalini (Coleoptera: Scarabaeidae: Dynastinae)

**DOI:** 10.1101/2025.09.18.677223

**Authors:** Wonseok Choi, Adrian Troya

## Abstract

The endophallus is the intromittent, membranous structure inside the aedeagus. It provides unique information for distinguishing taxa: species and genera. In addition to taxonomy, this organ is also useful to get insights about the evolution of the studied taxa, thus informing their classification. Despite the extensive reports on the structure and function of insect genitalia, the endophallus has received less attention than the sclerotized structures of the aedeagus. Several genera in the Agaocephalini rhinoceros beetles can be classified in three distinct groups based on the morphology of their endophalli. By examining the genitals of about 60% of the currently recognized species in this morphologically heterogeneous tribe, we provide for the first time a general overview of this structure. Furthermore, based on existing differences between genera, we discuss about the potential implications for the internal tribal systematics, as currently accepted. Novel information about endophalli structures will shed light on the missing links to other dynastine tribes.

## Introduction

The genital structures have been widely used in taxonomy and systematics in Coleoptera (Blaisdell, 1939; Miller, 2001; Medina et al., 2013; Schilthuizen, 2016), and their relatively rapid divergence is especially convenient in distinguishing closely related species (Eberhard, 2010), but also for studying the classification of beetles above the species level in some families (Tschinkel & Doyen, 1980; Calder, 1990). The taxonomic utility of the multicomponent genital system, which has its basis on the comparison of intra-population versus inter-population variants (Cohn, 1994), has led to numerous discoveries driving the progress of morphological research in beetles. Because of this, the genital structures are extensively reported in the literature.

The scarab beetles (Scarabaeidae) show some of the most extreme cases of inter- and intrapopulation morphological disparities, usually polyphenic or allometric in nature (Moczek, 2002; Rowland, 2005; Warren et al, 2014), most cases of which are well-exemplified in the head armature of the long-horned beetles (Dynastinae). This phenotypic plasticity usually obscures taxonomic differentiation (Moczek, 2002). Besides the examination of external body structures, and in the absence of alternative sources of evidence, for example, DNA, ecology, behavior, the morphology of the genitalia, also called terminalia (see, for example, Cristóvão & Vaz-de-Mello, 2021) can be of valuable support to the taxonomist. Most species of dynastines display species-specific genital structures which are informative for their diagnoses (Ratcliffe, 2021). The morphological features of the male’s genitalia, or aedeagus, like the parameres, which are involved in tactile stimulations (Düngelhoef & Schmitt, 2010), the phallobase, and the endophallus, which is a sperm-transferring device, are the main structures of interest.

The external sclerotized structure of the aedeagus is composed of a bilobed tegmen, formed by two parameres (Fig. 2A, B), and a phallobase (Fig. 1B) (Cristóvão & Vaz-de-Mello, 2021; Scholtz, 1990). Whereas the endophallus, which is a membranous sac encased inside the aedeagus, specifically within the median lobe, is composed of two sclerotized temones, a lobe (or lobes), and a number of spine-like endophallites (Medina et al., 2013; Génier, 2019; Cristóvão & Vaz-de-Mello, 2021). In an evolutionary context the complexity of these structures could be the result of cryptic female choice (Roig-Alsina, 1993), acting as an important sexual selection mechanism (Sloan & Simmons, 2019). Contrary to the well-studied, conspicuous sclerotized structures of the aedeagus, the endophallus has received less attention, likely due to the difficulty of dissecting and accessing this complex, fragile intromittent organ. Recently, Cristóvão & Vaz-de-Mello (2021) provided a revised glossary of terms for the terminalia of Scarabaeoidea, including the structures of the endophallus, which was certainly needed.

**Figure 1.**
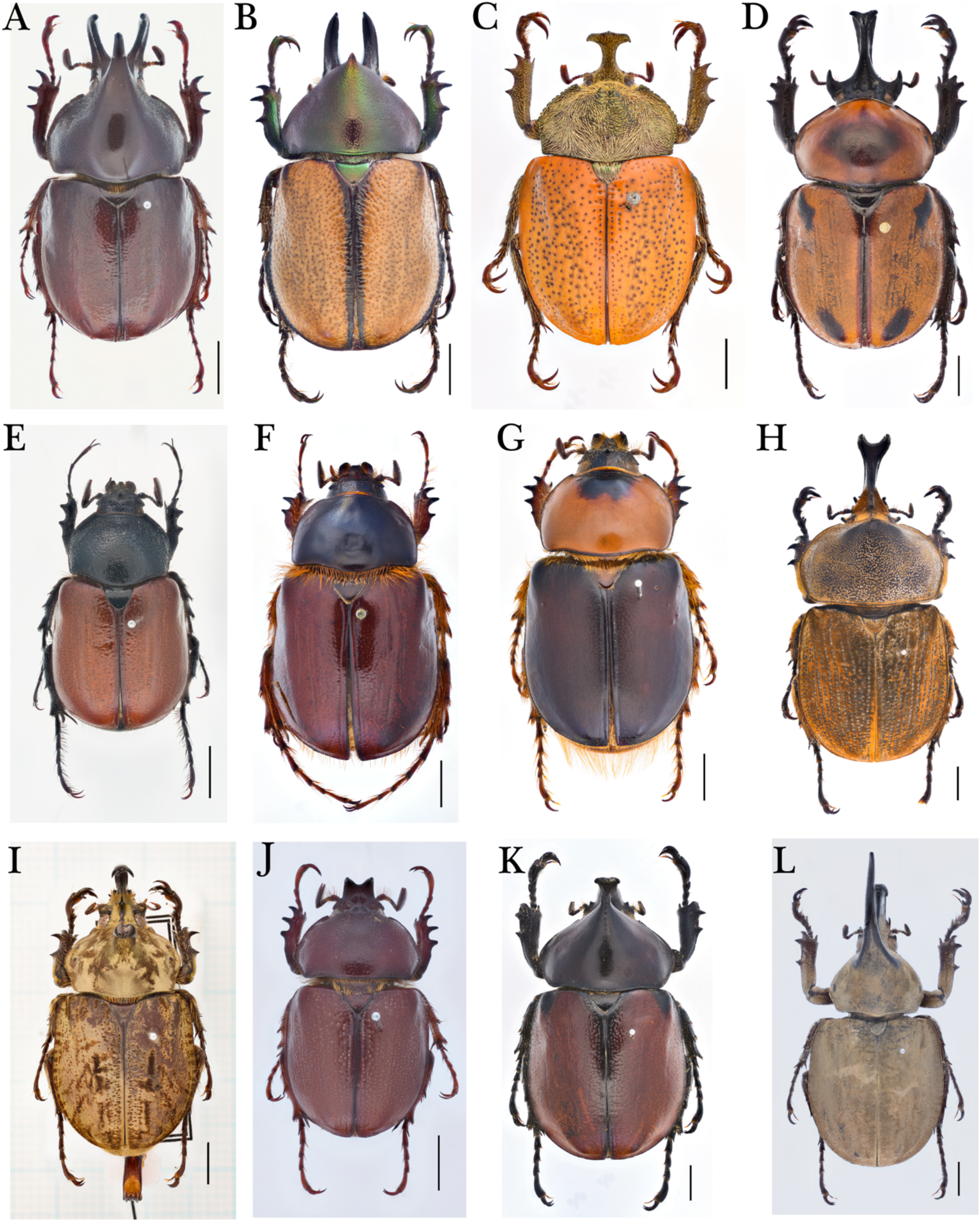
A sample of the male taxonomic diversity of Agaocephalini rhinoceros beetles whose endophalli were examined in this study. Each image represents the type species of its respective genus. (A) *Aegopsis bolboceridus* (Thomson); (B) *Agaocephala cornigera* Serville; C. *Antodon goryi* (Laporte); (D) *Brachysiderus quadrimaculatus* Waterhouse; (E) *Colacus bicolor* Ohaus; (F) *Democrates burmeisteri* Reiche; (G) *Gnathogolofa bicolor* (Ohaus); (H) *Horridocalia delislei* Endrödi; (I) *Lycomedes reichei* Brême; (J) *Minisiderus minicola* (Ohaus); (K) *Mitracephala humboldti* Thomson; (L) *Spodistes mniszechi* (Thomson). Scale bars: 5 mm.

**Figure 2.**
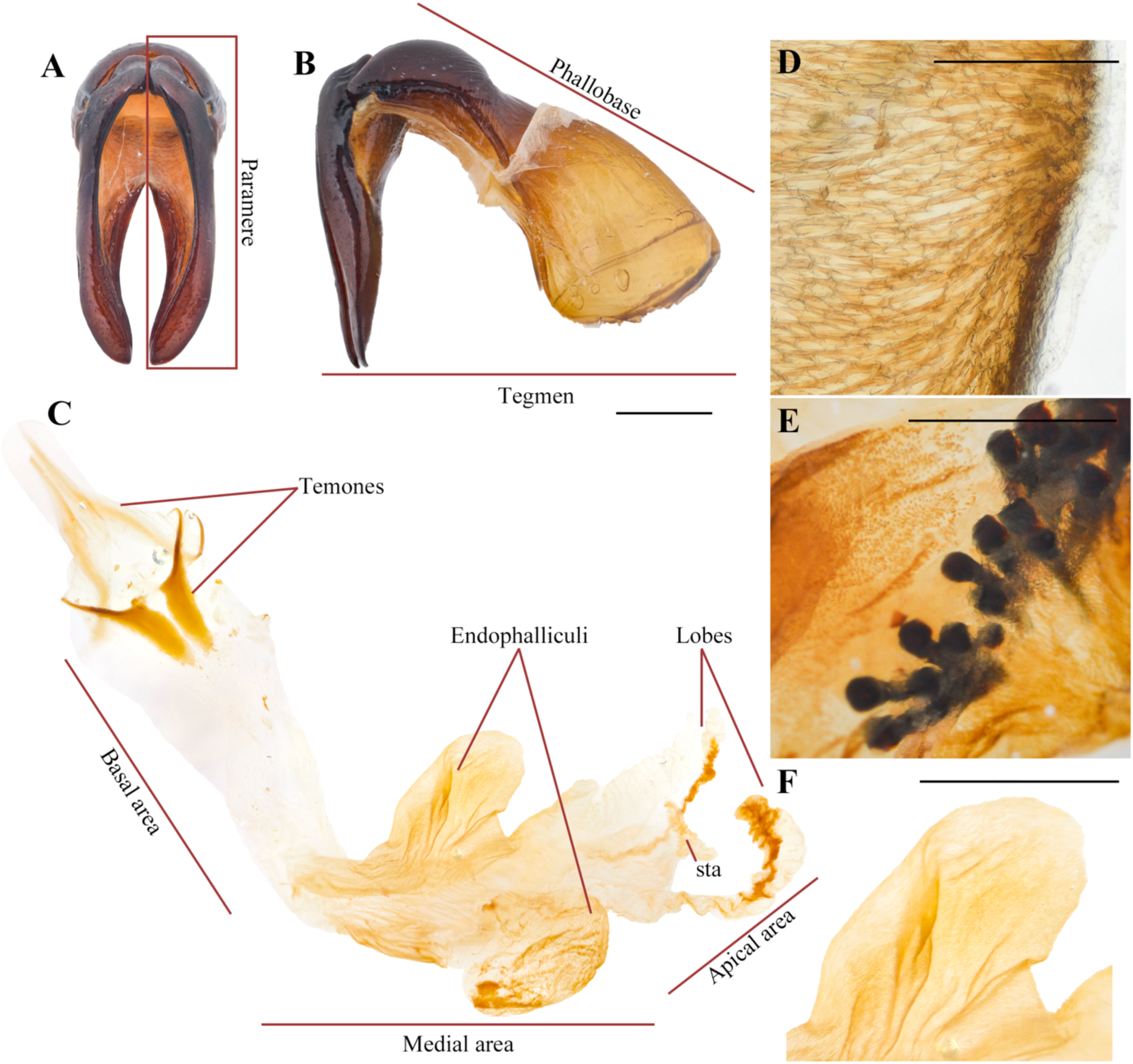
Main genital dynastine structures referred in this study. (A) Aedeagus, dorsal view; (B) Aedeagus, lateral view; (C) Endophallus; (D) Type I raspulae; (E) Type II raspulae; (F) Endophalliculus. **sta**: sperm transfer apparatus. Scale bars. C, E, F: 1 mm; D: 100 μm.

The genitalia of insects show a remarkable diversity due to selection mechanisms that have not been fully elucidated (Hosken & Stockley, 2004; Hunt, *et al*., 2009; Sasakawa, 2022). Nevertheless, due to its “rapid” evolution (Hosken & Stockley, 2004), genital characters are sometimes omitted in phylogenetic analyses, because genital traits may not be informative. Some studies, however, are proving otherwise, see for example, Song & Bucheli (2010); Zhou et al. (2020); Fang et al. (2023); Fernández-Campos et al. (2024).

The endophallus morphology is nowadays more understood than in the past and is increasingly used in taxonomic treatments of some dynastine groups. Morón (1995), for example, reported on features of the endophallites of the *Golofa* Hope of Mexico; Rowland (2003) suggested the morphology of the raspulae (a type of endophallite) may be a unique diagnostic trait in the genus *Xylotrupes* (Linnaeus); Neita-Moreno & Ratcliffe (2017) and Neita-Moreno (2021) provided details of endophallic traits in a review of *Tomarus* Erichson from Argentina, Chile, and Uruguay, and of *Cyclocephala* Dejean from Colombia, respectively.

In the present contribution we provide the first comparative overview of the endophallus morphology of selected species representing all the component genera in the dynastine tribe Agaocephalini. This is a small group of about 60 species distributed in the Neotropics, namely Central America and South America. This tribe is hard to diagnose because of the heterogeneity of external morphological characters of the body of its component genera (Ratcliffe, 2003). Endrödi (1985), for example, pointed out that *Democrates* Burmeister and *Colacus* Ohaus are closely related to the Cyclocephalini. More recently, Sobral (2023) suggested to move those genera to the Pentodontini. Based on our observations we here also comment on their suggested changes.

## Material and methods

### Specimen preparation

General procedures and terminology followed Cristóvão & Vaz-de-Mello (2021). Selected specimens were softened by immersion in hot water for 10-20 minutes. The entire abdomen was removed, and its contents were softened including the aedeagus which was extracted as described in Ratcliffe (2021). The removed tissues were immersed in 10% KOH solution at room temperature until connective tissues were digested, in average about 6 hours. Older specimens or badly preserved needed longer digesting time up to 24 hours. After internal tissues were digested and softened, whole tissues including genitalia were transferred to distilled water. Membranous structures were inflated due to osmosis, and spontaneously a part of the endophallus everted out of the phallobase. The endophallus was extracted by gently pulling out the temones and median lobe, then placed in alcohol. If the endophallus remained solid or was difficult to observe due to opaque fragments, it was reimmersed in a 10% KOH solution for 24 hours. Each piece was washed in 70% ethyl alcohol and stored in glycerol for posterior examination.

### Terminology

We divided the endophallus in the following three areas, so as to facilitate the recognition of specific structures: 1. A basal area where the temones are embedded; 2. A medial area where most endophallites and endophalliculi are placed; 3. An apical area bearing the lobes and the sperm transfer apparatus (Medina et al., 2013; Fig. 2C).

The term ‘endophalliculus’ (singular) and ‘endophalliculi’ (plural), here first proposed, refers to the relatively small sacs usually covered with minute type I raspulae (Fig. 2D), which extend from the main endophallus body (Fig. 2F). This term is a neologism, and is composed of the words ‘endophallus’, and the masculine form of the Latin diminutive suffix ‘culus’, meaning ‘little’. The other narrow, long, and tube-like form of extended structures from the main body is referred to ‘lobe’, following the usage of Genier and Morreto (2017).

The term ‘raspula’ (singular) or ‘raspulae’ (plural), follows Cristóvão & Vaz-de-Mello (2021), and describes two types of endophallites: Type I, which are densely grouped, conical-shaped setae, typically placed in the medial area, and around the endophalliculi (Fig. 2D); and Type II, which are spine- or tooth-shaped setae are most often placed on the endophalliculi and on the main endophallus body (Fig. 2E).

Finally, the term ‘columna’ (singular), or ‘columnae’ (plural), here first proposed, refers to the usually longest section of each temone, which is connected to its respective arm. Both the arm and the columna form a single structure. Here, the aim of naming a new term is to facilitate the distinction and description of the examined morphological feature.

### Specimen observation

The prepared endophalli were mounted in 2.5% carboxymethyl cellulose (Samchun Co. Ltd., product # C0292) solution on microscope slides. Some endophalli were hard to observe due to transparency, thus they were stained with 10% Nigrosin (Junsei Co., Ltd., product # 53075-1210) aqueous solution.

To examine the structures of inflated endophalli, 2.5% carboxymethyl cellulose solution was injected through the phallobase using a 15 g oral zonde needle. The inflated endophalli were photographed immediately and stored in glycerol.

### Imaging

All specimens were examined with a Nikon SMZ645 stereoscope and Eclipse 50i microscope, then photographed with a Nikon D5200 coupled with a Tamron 90 mm Macro lens and Laowa 25mm f/2.8 Ultra 2.5-5X lens mounted on a Wemacro macro rail. The resulting images were stacked using Zerene Stacker (Zerene Systems LLC) and retouched using Adobe Photoshop. All images by the first author.

### Material examined

In order to compare homologous structures, besides the below selected Agaocephalini species, we also included members of the tribes Cyclocephalini, Dynastini, Oryctini, Pentodontini, and Phileurini (Table 1). The specimens were identified by the first author using the taxonomic treatments of Endrödi (1970, 1985), Ratcliffe (2003), Ratcliffe et al (2020), Sobral (2023), as well as by examining specimens in the following collections: Wonseok Choi Personal Collection – WCPC; Natural History Museum London – NHM; Muséum National d’Histoire Naturelle – MNHN. The material, examined specimens and preparations, are preserved in the WCPC collection and is freely available for research upon request.

**Table 1.**
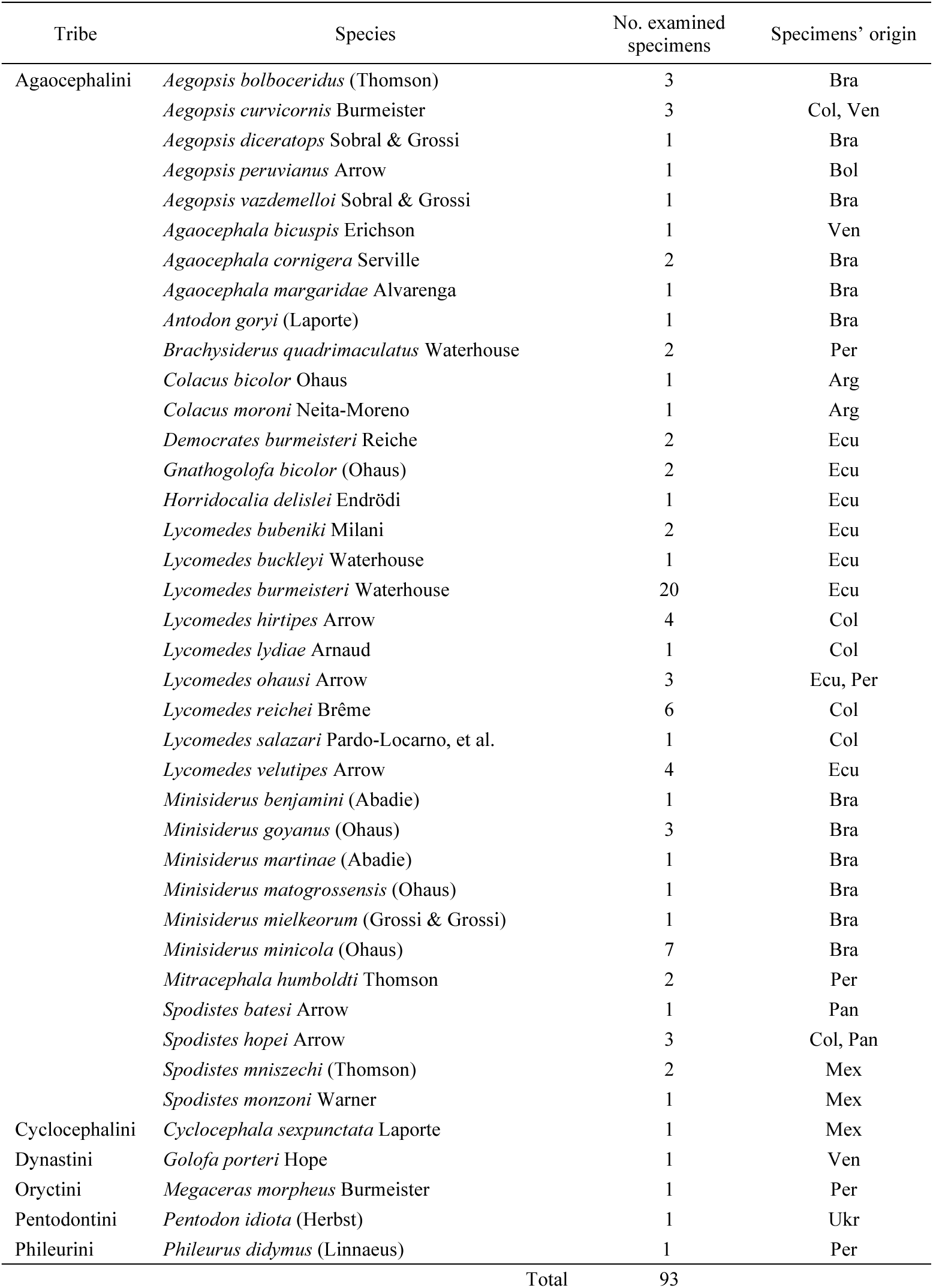
Material examined in this study. Abbreviations. Arg: Argentina, Bol: Bolivia, Bra: Brazil, Col: Colombia, Ecu: Ecuador, Mex: Mexico, Pan: Panama, Per: Peru, Ukr: Ukraine, Ven: Venezuela.

## Results

### General overview of endophallus morphology of the Dynastinae

The morphology of the endophallus in examined dynastines is overall similar with respect to the presence of two symmetric, elongated, and usually Y-shaped temones forming a ring-like structure (Fig. 3A), and a relatively long main sac bearing one or more endophalliculi (Fig. 3B). While the basal and apical area show little variations in shape, the medial area varies broadly in terms of the number and shape of lobes, endophalliculi, and endophallites. For example, *Golofa porteri* (Dynastini) shows four endophalliculi, three of which bear endophallites and two are distally connected by narrow lobes (Fig. 3B); *Cyclocephala sexpunctata* (Cyclocephalini; Fig. 3C) bears a single, uniformly sized endophalliculus and an elongated narrow lobe; *Pentodon idiota* (Pentodontini; Fig. 3D) bears four lobes; *Megaceras morpheus* (Oryctini; Fig. 3E) shows an enlarged endophalliculus; *Phileurus didymus* (Phileurini; Fig. 3F) bears an enlarged endophalliculus extended from the medial area and numerous lobes.

**Figure 3.**
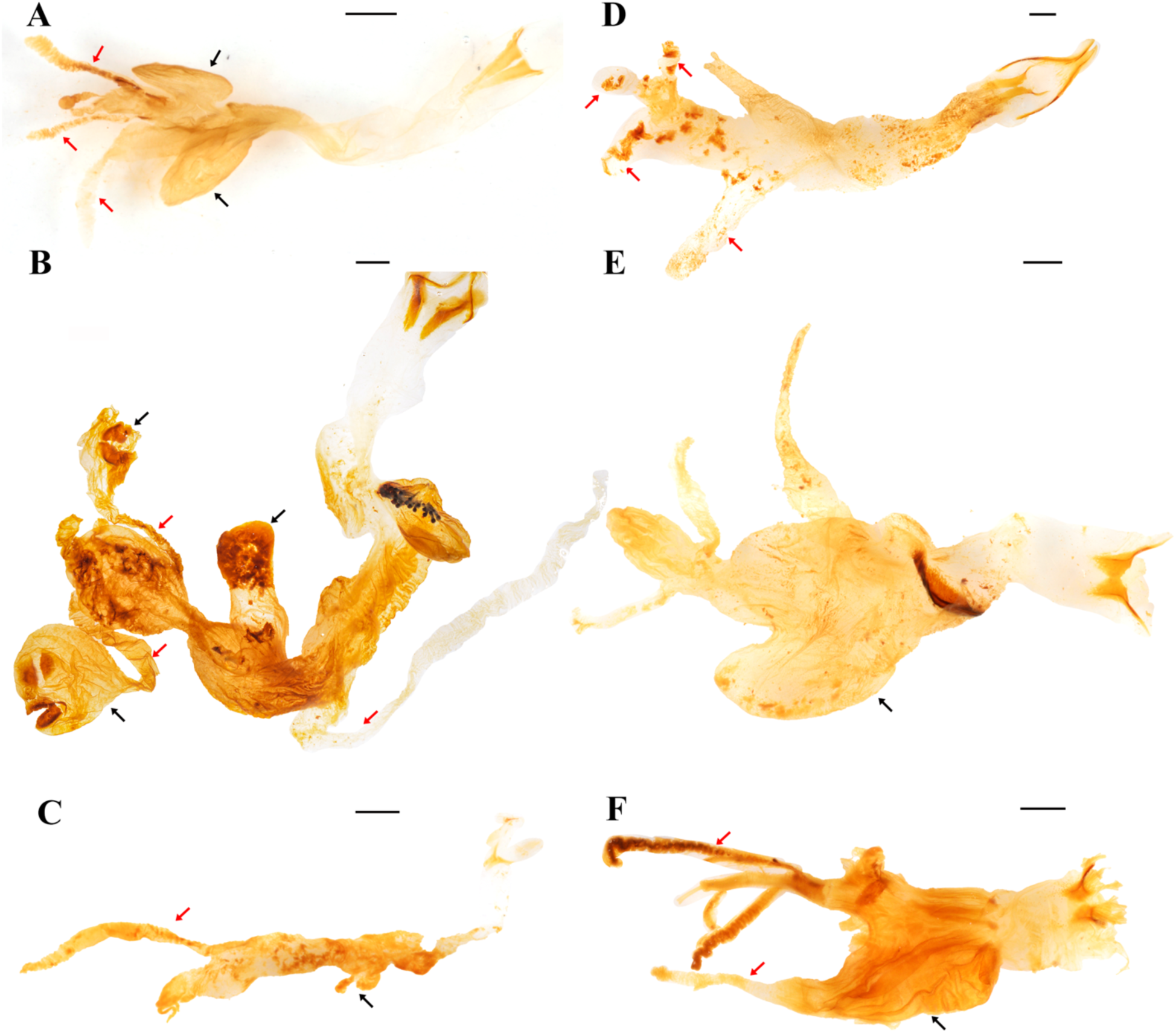
General view of the endophallus in some Dynastinae tribes. (A) *Lycomedes reichei* Brême (Agaocephalini); (B) *Golofa porteri* Hope (Dynastini); (C) *Cyclocephala sexpunctata* Laporte (Cyclocephalini); (D) *Pentodon idiota* (Herbst) (Pentodontini); (E) *Megaceras morpheus* Burmeister (Oryctini); (F) *Phileurus didymus* (Linnaeus) (Phileurini). Black arrows point to the endophalliculi; red arrows point to lobes. Scale bars: 1 mm.

Endophallites may be present or absent depending on each species, and their shape and number vary accordingly. For example, *Cy. sexpunctata* (Fig. 3C), *Pe. idiota* (Fig. 3D), *Ph. didymus* (Phileurini) (Fig. 3F) are not armed with prominent endophallites. Type I raspulae with dense conical setae (Fig. 2D, see also Fig. 5A in Binaghi et al., 1969) are present in *Ph. didymus*, and type II toothed raspulae (sensu Cristóvão & Vaz-de-Mello, 2021) are present in *Go. porteri* (Fig. 4A). Additionally, endophallites shaped as plates as in *Go. porteri* (Fig. 4B) and *Me. morpheus* (Fig. 4C), or as conical plugs as in several Agaocephlini species, including *Mit. humboldti*, *Ag. cornigera* (Fig. 4D), can be distinguished. The apical area of all observed endophalli shows a similar structure with two lobes, and a short, narrow, tubular structure at the vertex of the main endophallus body (Fig. 4E).

**Figure 4.**
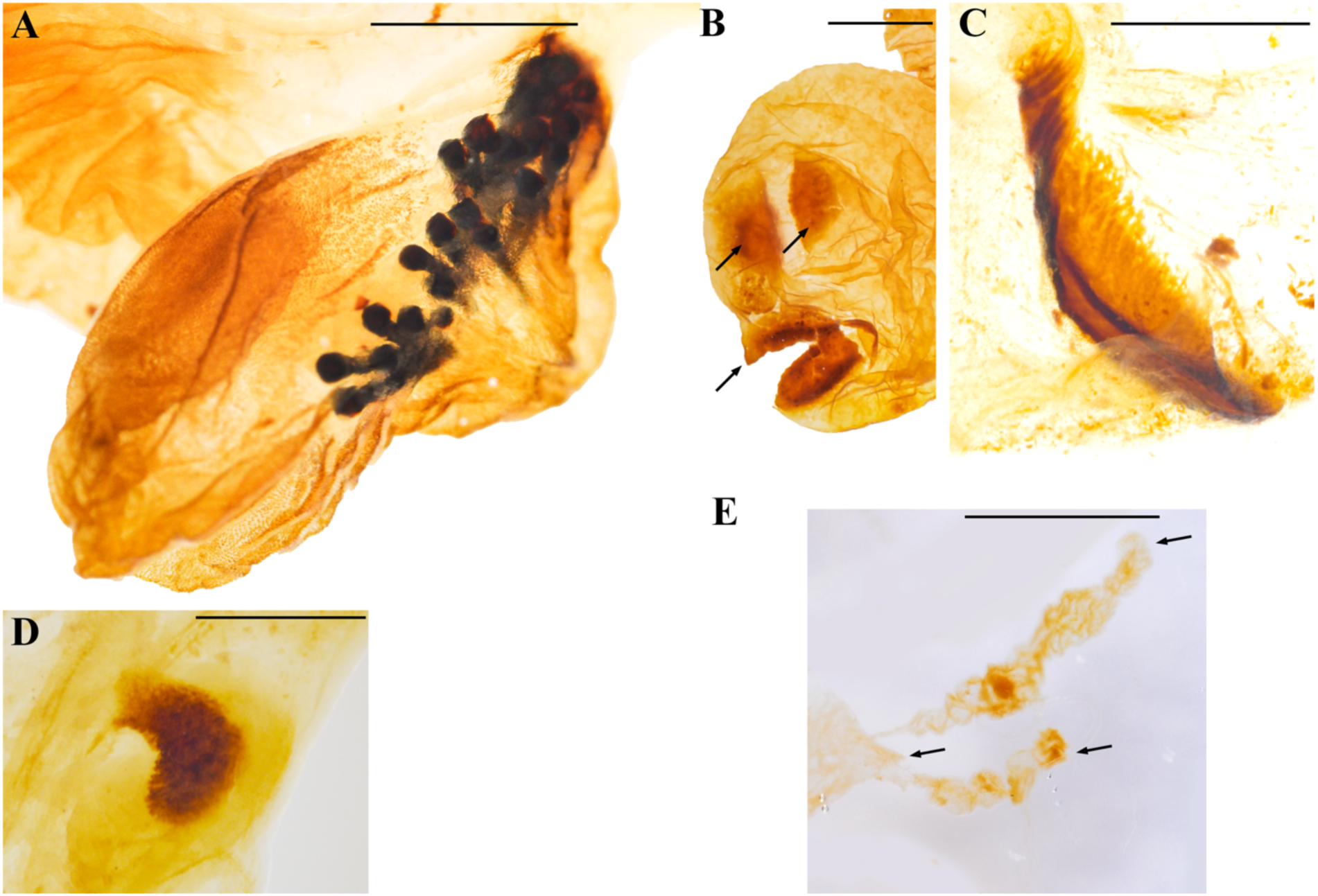
Close ups of some morphological features of the endophallus in Dynastinae. (A) Type II raspulae (*Golofa porteri* Hope); (B) Plate-shaped endophallites (black arrows) (*G. porteri*); (C) Plate-shaped endophallite (*Megaceras morpheus* Burmeister); (D) Endophallite of *Mitacephala humboldti* Thomson; (E) Apex of endophallus bearing two lobes and a short tubular structure at the vertex (black arrows), (*Lycomedes burmeisteri* Waterhouse). Scale bars: 1 mm.

**Figure 5.**
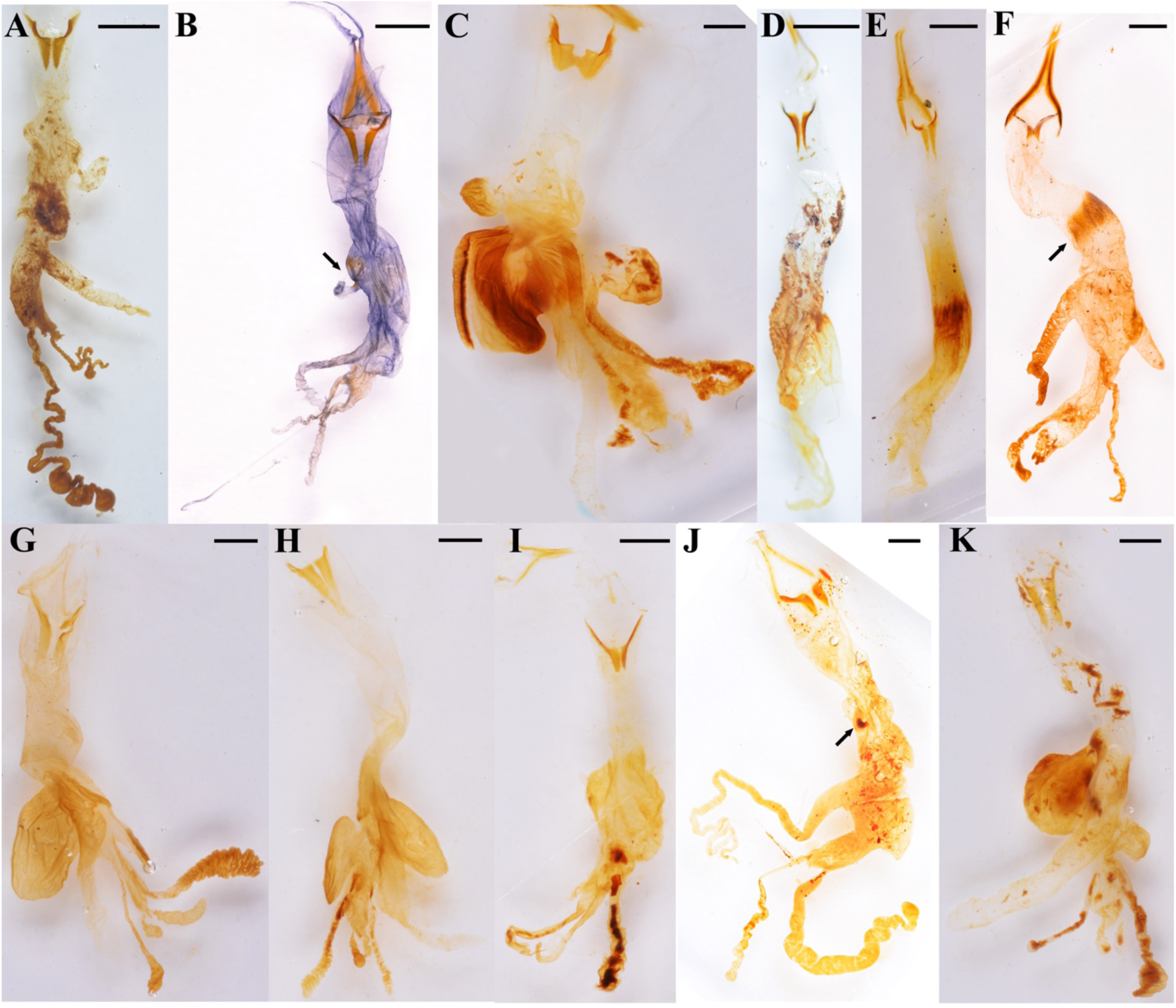
Endophallus of the Agaocephalini. The complete organ is shown for each genus, except *Antodon*, which was damaged while dissecting it. (A) *Aegopsis curvicornis* Burmeister; (B) *Agaocephala cornigera* Serville; (C) *Brachysiderus quadrimaculatus* Waterhouse; (D) *Colacus bicolor* Ohaus; (E) *Democrates burmeisteri* Reiche; (F) *Gnathogolofa bicolor* (Ohaus), black arrow points to seta-like hairy type I raspulae; (G) *Horridocalia delislei* Endrödi; (H) *Lycomedes reichei* Brême; (I) *Minisiderus matogrossensis* (Ohaus); (J) *Mitracephala humboldti* Thomson (black arrows points to endophallites at the connecting area between endophalliculus and the main endophallus body); (K) *Spodistes batesi* Arrow. Scale bars: 1mm.

### The endophallus of the Agaocephalini Basal area

The temones vary in size and shape among genera. We will refer here only to the pair of temones whose arms are in opposed direction to the other endophallic structures because in most cases the other pair was broken in the dissection process. In *Ae. curvicornis,* the length of the proximal region, this is, the columna of each temone is about twice as long as its base (Fig. 5A), whereas in *Ag. cornigera* (Fig. 5B), *Co. bicolor* (Fig. 5D), *H. delislei* (Fig. 5G), *L. reichei* (Fig. 5H), and *S. batesi* (Fig. 5K), it is three times longer or more. The columnae of several *Lycomedes* species, including *L. hirtipes* and *L. reichei*, show acute and well-defined vertices (Fig. 5H). On the contrary, other species in the same genus exhibit sub-triangular temones, which are less acute and have round vertices (Fig. 6A). In *Min. matogrossensis*, the columna is about as long as wide (Fig. 5I). In *Gn. bicolor*, we could not compare said proportion because the columna is incomplete (Fig. 5F). The temonal arms in this species, as well as those of *Co. moroni* (Fig. 6B), are peculiar because there is no distinction between a differentiated arm and the base of the columna. Here, the arm is as broad as the base of the columna and runs continuous with it, forming a bull’s-like horn structure (Fig. 5F). In *Gn. bicolor* the arm is directed lateriad, while in *Co. moroni,* it is strongly curved (Fig. 6B). Interestingly, the temones of *(B) quadrimaculatus* are connected medially (Fig. 5C), this was observed only in this species.

**Figure 6.**
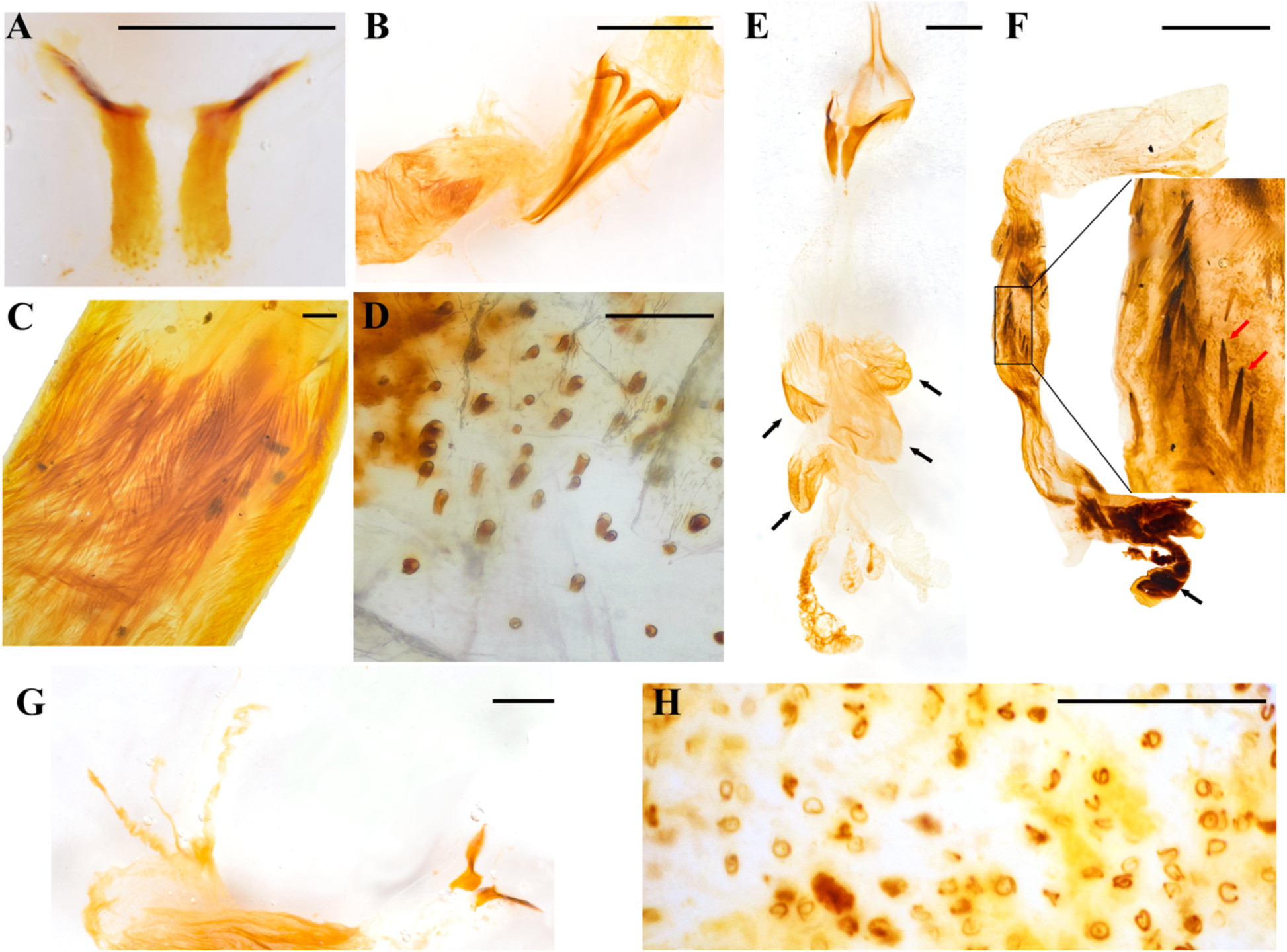
Close-up of some morphological features of agaocephaline endophalli. (A) Temones of *Lycomedes burmeisteri* Waterhouse; (B) Temones of *Colacus moroni* Neita-Moreno; (C) Hair-like type I raspulae at the endophallic medial area of *Democrates burmeisteri* Reiche; (D) Conical-shaped type I raspulae in *Lycomedes burmeisteri* Waterhouse; (E) Endophallus of *Lycomedes velutipes* Arrow (black arrows point at the endophalliculi); (F) Endophallus of *Antodon goryi* (Laporte) (the black arrow is showing the club-shaped lobe, and the red arrows indicate Type II raspulae); (G) Endophallus of *Aegopsis bolboceridus* (Thomson); (H) Ring-shaped endophallites of *Mitracephala humboldti* Thomson. Scale bars. A, B, C, E, F, G: 1mm; D, H: 100 μm.

### Medial area

A number of endophalliculi and lobes are extended throughout the medial area. *Lycomedes velutipes* has four endophalliculi (Fig. 6E), whereas the other species in that genus have three. *Horridocalia delislei* (Fig. 5G) and *S. batesi* (Fig. 5K) show two to three endophalliculi covered with type I conical raspulae, as well as the presence of one long lobe which is similar to that of *Lycomedes. Democrates burmeisteri* (Fig. 5E) and *Gn. bicolor* (Fig. 5F) show a long distinctive band of hair-like type I raspulae (Fig. 6C), while *L. bubeniki*, *L. burmeisteri,* and *L. ohausi* show conical-shaped type I raspulae (Fig. 6D). The raspulae at the basal area are more sparse and thicker than that at the surface of the medial area.

*Antodon goryi* is unique among the Agaocephalini because it is the only species that has distinct type II raspulae, which are thick and spine-shaped (Fig. 6F). The endophallus of *An. goryi* has one endophalliculus. The endophalli of *Ae. bolboceridus* and *Ae. curvicornis* exhibit clear differences. The latter has one endophalliculus and one lobe (Fig. 5A), whereas *Ae. bolboceridus* has an enlarged medial area covered with dense type I raspulae, with the lobes extending to the apical area (Fig. 6G). The endophalliculus of *Brachysiderus quadrimaculatus* as well as that of all species in *Lycomedes* is similar in shape and length (Fig. 6D), but with distinct type I raspulae. Here, part of the endophalliculus is covered with a band of dense setae, and the rest is conical-shaped, a feature shared by all species in *Lycomedes* (Fig. 5C).

The endophallic medial area of *Ag. cornigera* (Fig. 5B) and *Mit. humboldti* (Fig. 5J) has one endophalliculus; the connecting area between it and the main body is blocked by a single knob-like endophallite (Figs. 5B, 5J). *Agaocephala Cornigera* has a patch of type I raspulae, which are similarly placed, this is, at the juxtaposition of the endophallite (Figs. 5B, 5I). However, *Min. minicola* does not have endophallites except for minute surface-covering type I raspulae. Interestingly, we noted that the endophalli of some specimens of *Mit. humboldti* and *Min. goyanus* have a small, transparent lobe, where ring-shaped, small endophallites are embedded (Fig. 6H).

### Apical area

All examined Dynastine species exhibited a similar configuration of features in the apical area. Two lobes and a narrow tubular structure at the vertex are observed. These lobes vary from club-shaped, as in *An. goryi* (Fig. 6F), to long and twisted, as in *Ae. curvicornis* (Fig. 5A), to simple and long, as in the remaining species.

## Discussion

### Heterogeneity of Agaocephaline endophalli and its implications for the internal tribal classification

Our observations underscore considerable morphological diversity of the endophallus in the examined Agaocephaline species, particularly in the form and composition of sclerites. This heterogeneity contrasts with the more conservative, but diverse configurations reported in Scarabaeinae lineages. In Onthophagini, for example, the endophallic morphology is highly informative due to its low homoplasy, which means that endophallic characters likely correlate with the observed phylogenetic relationships among the genera and species in that tribe (Tarasov and Solodovnikov, 2011). Comparable patterns of variability have been documented in Ateuchini (Kohlmann and Solis, 2009), Dichotomini (Montoya-Molina and Vaz-de-Mello, 2021), Sericini (Ahrens, 2006), Phaneini: *Bolbites onitoides* Harold, 1868 (Cupello et al., 2022), Trichiini (Lis et al., 2008), and Deltochilini: *Canthon* Hoffmannsegg, 1817. In this regard, Medina et al. (2013) demonstrated that, despite the high variation of shared traits among several dung beetle taxa, the endophallus morphology is generally stable so as to characterize broader phylogenetic patterns across the subfamily. In contrast, at the intergeneric level, our observations reveal that Agaocephalini exhibits an even greater degree of variation reflected in the presence or absence of sclerites, as well as in their shape, than what has been published in Scarabaeinae.

The most noticeable difference among the examined Agaocephalini species is the type and presence of endophallites. Hairy, seta-like type I raspulae at the medial area are found in *Co. moroni* (Fig. 6B), *D. burmeisteri* (Fig. 6C), and *Gn. bicolor* (Fig. 5F). Distinct type II raspulae are only found in *An. goryi* (Fig. 6F). *Mitracephala humboldti* and *Ag. cornigera* have endophallites or raspulae at the connecting area between the main body and an endophalliculus (Figs. 5B and 5J). The remaining species listed in the Table 2, except *Lycomedes enigmaticus, L. ramosus*, *Minisiderus bertolossiorum*, *Min. elyanae*, and *Min. paranensis*, which were not examined in this work, have only minute-sized, type I raspulae (Figs. 2D, 5F).

**Table 2.**
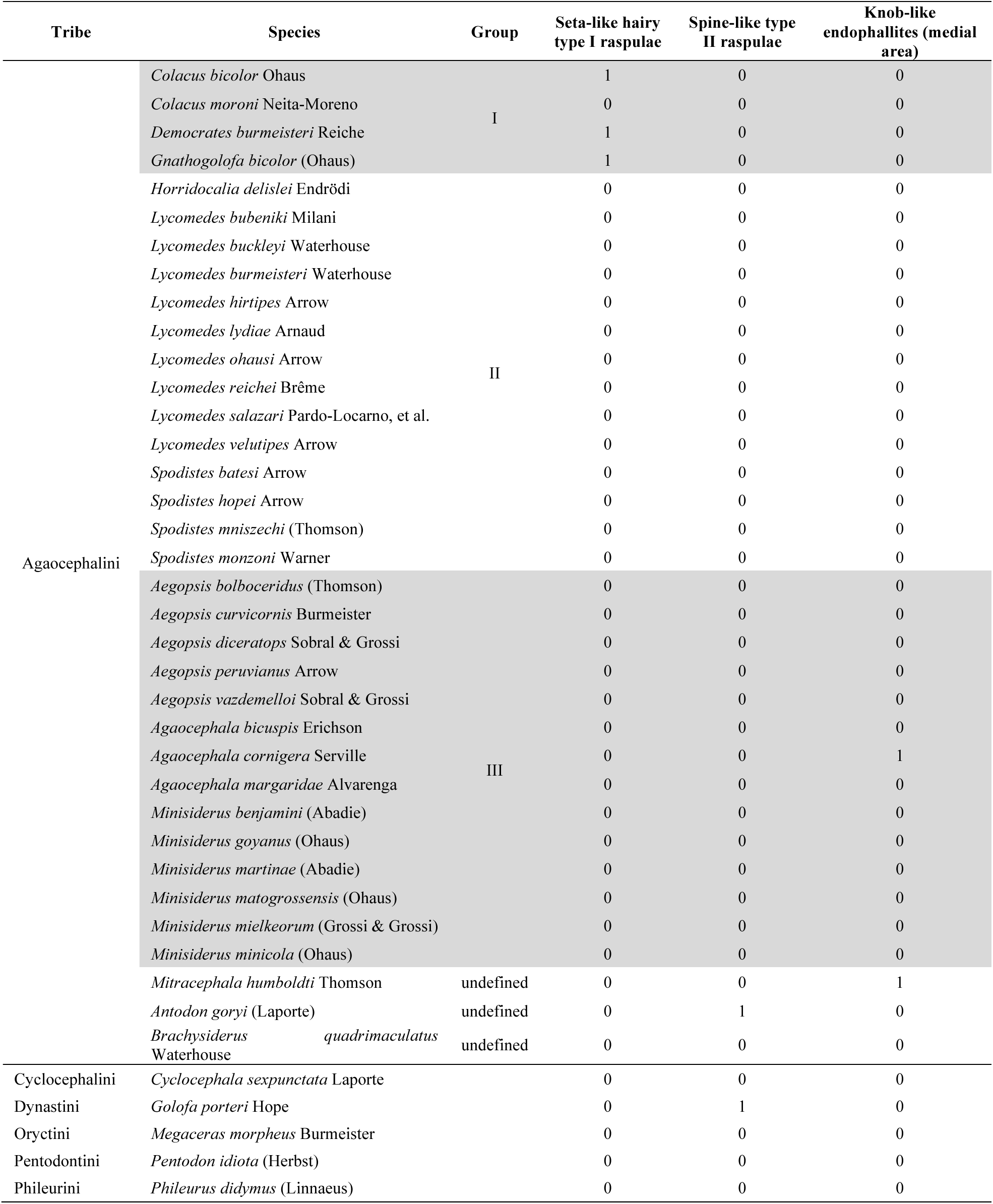
Differences in certain endophallite features along all examined species. 1= present; 0 = absent.

The morphological characters reflecting this variation, however, are apparently not shared and derived along all the lineages that compose the tribe, as currently defined. This not only raises questions about the phylogenetic relationships among the currently known genera in the tribe, but also about the functional and evolutionary significance of endophallus architecture.

The observed differences in general endophallus morphology along our sample of agaocephaline species reinforce our idea of a paraphyletic tribe. The presence of specialized structures, such as type I and II raspulae, as well as various forms of endophalliculi, show a degree of divergence which we cannot explain from an evolutionary perspective based on the current sample size (93 individuals), even though these represent a relatively broad geographic range in South America (Table 1; Supplementary Material, available online). A future phylogenetic analysis using a scoring matrix including endophallic- and other external morphological traits, is needed to dig deeper into the relationships among genera.

It remains to be tested whether some traits like the number of lobes or shape of the raspulae are synapomorphic along the tribe, as currently defined. However, the observed features appear to cluster in particular lineages, for example, narrow and small lobes (Figs. 5D-5F) are present only in *Colacus*, *Democrates* and *Gnathogolofa*, here classified under group I (Table 2), whereas a broad and large endophalliculus (Figs. 5G, 5H, 5K) is present in *Horridocalia*, *Lycomedes*, and *Spodistes*, here classified under group II (Table 2). Group III (Table 2), on the other hand, shows a modestly extended endophalliculi as in *Minisiderus* (Fig. 5I), a minute endophalliculus connected to the main endophallus body, as in *Agaocephala* (Fig. 5B), or with both types of endophalliculi as in *Aegopsis* (Fig. 5A). In addition, long and often twisted, tube-like lobes are present in the three genera comprising group III (Figs. 5A-B, 5I). As of the remaining three genera, *Antodon*, *Brachysiderus*, and *Mitracephala*, we could not find enough evidence, either from endophallic structures or from external morphological characters, to justify their placement within any of the aforementioned groups. The raspulae of *An. goryi* and *Br. quadrimaculatus* are differently shaped, with the first showing type II (Fig. 6F), while in the second the raspulae form a dense band of seta-like type I (Fig. 5C). Although *Mit. humboldti* (Fig. 5J) shares some endophallic traits with species in group III, it differs with all of them by a number of external body traits, most notably a single, thick cephalic horn and thickened protarsi, while all species of group III show two cephalic horns and thinner protarsi. *Mitracephala humboldti* has been collected in montane forests of the northern Ecuadorian Andes, Peru, and northern Yungas of Bolivia, whereas most species in group III have been recorded from the Brazilian Cerrado, except for *Ae. curvicornis*, *Ag. bicuspis*, and *Ag. margaridae*, which are known only from Andean or Amazonian sites in Venezuela, Colombia, and Northern Brazil.

The three groups are also supported by patterns in external morphology. Genera in group I lack a pronotal horn, have a tubercle on the frons (Figs. 1E-G), symmetric claws, broad and leaf-like mandibles, and the body has a general matte surface. Also, the parameres of group I genera, as seen in dorsal view, are slender and elongated, with short apical setae (Figs. 7A-C). Species in Group II have prominent cephalic and pronotal horns (Figs. 1H, 1I, 1L), asymmetric claws, mandibles with two to three teeth, and the body has a tomentose surface (Fig. 1L). The parameres of group II genera, as seen in dorsal view, are less elongated than those of group I genera, and are broadened distally, without apical setae (Figs. 7D-F).

**Figure 7.**
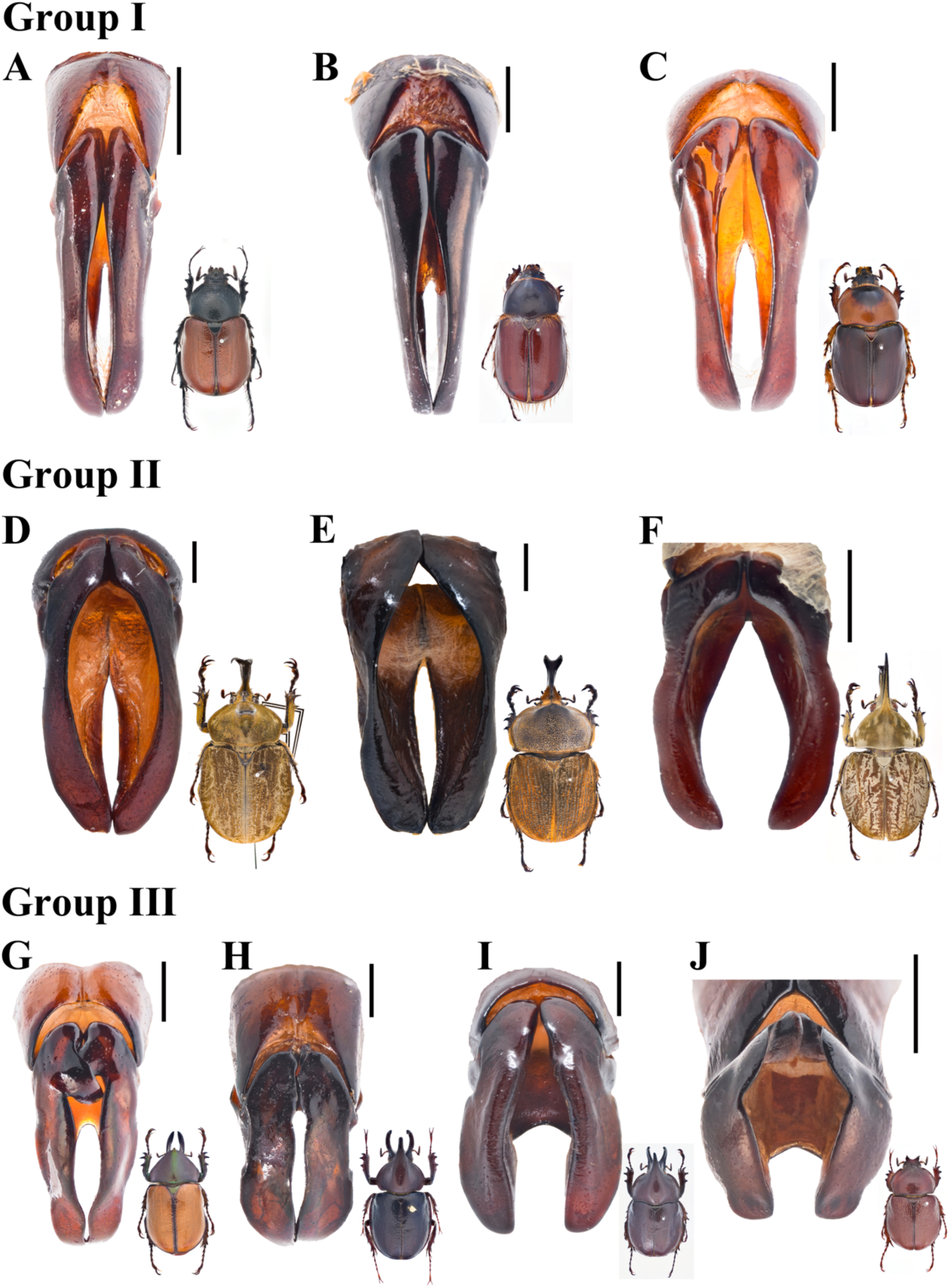
Parameres (dorsal view) of the three here recognized agaocephaline groups. Group I: (A) *Colacus bicolor* Ohaus; (B) *Democrates burmeisteri* Reiche; (C) *Gnathogolofa bicolor* (Ohaus). Group II: (D) *Horridocalia delislei* Endrödi; (E) *Lycomedes burmeisteri* Waterhouse; (F) *Spodistes grandis* Sternberg. Group III: (G) *Agaocephala cornigera* Serville. (H) *Aegopsis curvicornis* Burmeister. (I) *Aegopsis bolboceridus* (Thomson). (J) *Minisiderus mielkeorum.* (Grossi & Grossi). Images of adults not to scale. Scale bars: 1 mm.

The shared traits among genera in group III are not as evident as in the previous groups. For example, species in *Aegopsis* and *Agaocephala* bear two cephalic horns, while in *Minisiderus* the cephalic horn forms a single trunk proximally but diverges in two prominent teeth apically. (Fig. 1J). *Minisiderus* bear a knob-like tubercle (Fig. 1J) on the pronotum, while in *Aegopsis* and *Agaocephala* this tubercle is protruding anteriad and is larger than in *Minisiderus* (Figs. 1A-B), with the exception of *Ag. bicuspis*, *Ag. duponti*, and *Ag. inermicollis*. The parameres in these genera, in dorsal view, are usually asymmetric and somewhat rectangular (Figs. 7G-I), except for those in *Minisiderus* which are broadened medially (Fig. 7J). The protarsal claws are symmetric in most species within this group, whereas only in *Ag. cornigera*, *Ag. mannerheimi,* and *Ag. urus,* these claws are asymmetric.

The configuration of these traits may be the result of selective pressures, perhaps linked to reproductive isolation or mating mechanisms like the structural complexity of endophallic architecture, which may have shaped the evolution of those lineages (Rönn et al, 2007). For example, the length of the endophallus has been associated with that of the female ovipositor in some cerambycid beetles of the tribes Trachyderini and Torneutini (Long & Tong, 2025). The exclusive occurrence of type II raspulae, in *An. goryi* points to a lineage-specific innovation, whereas the general reduction or complete absence of endophallites in other species, such as *Min. minicola*, a member of group III, may represent a secondary loss, given that these structures are otherwise present in all remaining members of the group. Further evidence is needed, however, to confirm or refute our hypothesis that the aforementioned genera constitute natural clades.

Some endophallic traits correlate geographically, thus reflecting possible divergence among certain lineages. We noticed this in two species of the genus *Aegopsis* whose distribution is clearly distinct. *Aegopsis curvicornis* is mostly Andean with its populations inhabiting mountainous regions of Trinidad & Tobago, Panama, Colombia and Ecuador (Endrödi, 1985, Choi et al., 2023). The endophallus of this species has a narrow body, one endophalliculus, and multiple long lobes (Fig. 5A). The other species, *Ae. bolboceridus*, inhabits the Brazilian Cerrado and the southern region of the Atlantic Forest (Endrödi, 1985, Choi et al., 2023). The endophallus of this species is mostly broad with narrow lobes (Fig. 5G). In contrast to their internal endophallus morphology, these two species are very similar externally, differing only in the number of tibial teeth, and in the form and size of their cephalic horns. All other species of *Aegopsis* that are distributed from Amazonian Peru to mid-southern Brazil, exhibited similar endophalli to that of *Ae. bolboceridus*.

### Asymmetry of temones

Asymmetry in genital structures is a common trait among insects (Schilthuizen, 2013). Although asymmetric parameres and their evolutionary implications have been examined, asymmetry of endophallic structures in Dynastinae has rarely been investigated (Breeschoten et al., 2013). In this study, the temones of all the Dynastinae examined were symmetric, except for that of *D. burmeisteri*. As far as we know, this is the first report of temonal asymmetry in an insect taxon. Whether this is an isolated case being the product of, for example, a random mutation generating a malformation, or it is in fact a trait of *D. burmeisteri*, it remains to be tested because we only examined a single individual of this rarely found species.

In a number of insect species, females nonrandomly distribute sperm into multiple spermathecae during post-copulatory selection, a phenomenon known as cryptic female choice (Birkhead, 1998; Hellriegel & Bernasconi, 2000; House et al., 2016). Temones may influence the directionality of the endophallus, possibly resulting in asymmetric sperm delivery. Further investigation is required to uncover the role of temones and the consequences of asymmetry in the context of female choice.

### Limitations of the endophallus as a tool in agaocephaline taxonomy

As noted before, the morphology of the examined endophalli allowed us to recognize three putative groups of genera. However, because the whole set of features may not be good synapomorphies, these may not be useful as a taxonomic tool for diagnosing the tribe, if its current taxonomic composition continues to hold after a proper phylogenetic study.

This is the first comprehensive examination of endophallus structures of a Dynastine tribe, however, without a broader understanding of endophallus morphology at the subfamily level, definitive conclusions about their taxonomic utility cannot yet be drawn. For example, the tribal placement of the three genera in group I remains uncertain. Endrödi (1970) noted that *Colacus*, *Democrates*, and *Gnathogolofa* might belong to Cyclocephalini, while Sobral et al. (2023) proposed transferring them to Pentodontini. Nevertheless, our current observations are insufficient to support either hypothesis, because the endophallus morphology of both tribes remains largely undocumented. Moreover, only 34 of the 57 known Agaocephalini species were examined in the current study. Broader taxonomic sampling and comparative analyses are needed to evaluate the utility of endophallus structures in the Dynastinae.

## Conclusion

This is the first attempt to scrutinize the complex morphological architecture of the internal intromittent organ of all currently accepted Agaocephaline genera. Although our sample size is still incomplete, we have shown that the observed characters are insufficient to formally propose a new classification for the tribe. Some features such as the number and shape of raspulae, endophalliculi, and lobes, proved useful for diagnosing some genus groups we have here identified. These groups, however, require deeper support, ideally integrating data from a comprehensive scoring matrix of both internal and external structures, alongside DNA evidence. Expanding the taxonomic sampling and combining these datasets will bring a higher resolution of the evolutionary relationships among genera, potentially leading to a formal reclassification of the of the Agaocephalini, which we suspect may not represent a natural group.

## Acknowledgements

AT thanks his family, especially Eli (his mom) for allowing resources at home to work in the manuscript.

## Notes

### Competing Interest Statement

The authors have declared no competing interest.

